# Critical assay parameters facilitating confident detection of expression changes, fusions and short variants in RNA isolated from tissue

**DOI:** 10.1101/2024.10.30.621018

**Authors:** Slawomir Kubik, Izabela Matyszczak, Nicholas Toda, Elia Magrinelli, Pierre-Yves Helleboid, Ewan Smith, Fedor Bezrukov, Chloe Ryder, Jonathan Lopez, Alexandre Harlé, Zhenyu Xu

## Abstract

Molecular assays based on next-generation sequencing of RNA can provide clinically relevant information by measuring gene expression levels, identifying gene fusions, aberrant transcript isoforms and detecting small variants. Nevertheless, achieving good performance and reliable result interpretation present a significant challenge.

In this work, we dissect the impact of various technical factors on the concurrent detection of several biomarker types using a hybridization enrichment-based targeted RNA sequencing applied to a cohort of more than one hundred samples derived from diverse solid tumors.

We demonstrate that several critical parameters inferred from the sequencing data should be controlled to interpret the results accurately. These include molecular coverage of reference genes as a proxy of RNA conversion, DNA content measure, and the extent of target molecular coverage. We summarize our findings with recommendations allowing maximum reliable information recovery from targeted RNA sequencing data.

## INTRODUCTION

Methods utilizing next-generation sequencing (NGS) became prevalent tools in tumor diagnosis and cancer management. While most assays traditionally rely on the analysis of DNA-derived data, specific insights are readily obtained using RNA as the input material.

Gene expression measurement may be used as proxy for gene amplification events, activating and silencing mutations or rearrangements. Expression profiles allow for determination of tumor heterogeneity^1^, prognosis^2^, prediction of treatment outcome^3^ or identification of new treatment options^4^. Targeted RNA-seq is particularly effective in detecting gene fusions, important biomarkers resulting from rearrangements that merge two otherwise distinct genes, leading to formation of hybrid transcripts^5^. Unlike DNA-based fusion detection, which requires deep sequencing of intronic breakpoints, RNA sequencing typically focuses on exon-exon junctions providing a direct and efficient screening approach. Detection of alternative transcript isoforms, such as exon 14-skipped MET^6^, is another example of alteration easily addressed by targeted RNA-seq. Both fusions and exon skipping events are associated with multiple tumor types^7,8^, and detection of an aberrant transcript may predispose the patient to targeted treatment^9–11^. Finally, RNA-seq data can also be used for direct detection of small variants^12^ which, together with information on allele-specific expression, allow insights into the tumor molecular profiles^13^. A single targeted RNA-seq dataset can thus serve as a versatile tool for obtaining a broad range of biologically and medically relevant information.

Correctly interpreting the RNA-seq results requires full awareness of the associated obstacles and limitations. For example, unlike DNA, RNA targets are present in the sample at non-equimolar amounts, leading to non-uniform performance across genes. Cost-efficient transcriptome interrogation requires enrichment of relevant targets, some of which may not be well expressed or may be subject to allelic imbalance^14^. The pitfalls are different when aiming for expression measurement versus those encountered in detecting of aberrant transcripts or small variants (small nucleotide variants - SNVs - and indels). Nevertheless, consistent recommendations on controlling overall data quality for the simultaneous detection of multiple types of biomarkers still need to be established.

Assays are often executed on material isolated from formaldehyde-fixed paraffin-embedded (FFPE) tissue samples. Such samples contain a variable amount of tumor cells and, consequently, a variable number of molecules bearing alterations, which impacts detection sensitivity. FFPE-derived nucleic acid is often fragmented and chemically degraded, which decreases the efficiency of converting RNA molecules into sequencing libraries^15,16^, causes sequence errors and increases noise^17,18^. Degradation of input material can be inferred from RNA size profiles and integrity metrics such as DV200 were shown to correlate with sequencing data quality^19^. However, fragmentation alone does not give information about the chemical damage of RNA. Reverse transcription is a step that can potentially elevate error levels^20,21^, diminishing the specificity of small variant calling in comparison to DNA-based assays^22,23^. Difficulty in discriminating between real chimeric transcripts and numerous sequencing artifacts requires stringent filtering. Consequently, weak fusion signal may be lost due to excessively conservative thresholds and fusions identified in the same dataset can differ depending on the fusion calling approach^24^. The presence of chimeric transcripts that do not result from genomic rearrangements further blurs the picture^25^. Finally, resident DNA in the specimen may bias the detection of all types of alterations^26^.

In this work, we extensively investigated the impact of input material features on the quality of RNA-derived targeted NGS data used for simultaneous measurement of gene expression, detection of gene fusions, alternative transcripts and small variants. We examined the effects RNA fragmentation, quality and quantity, and presence of DNA have on the test outcome and defined how it can be quantified in the data. We demonstrated that several metrics derived during the analyses can be used to evaluate the confidence of results and provide recommendations on how to interpret them for reliable detection of alterations.

## MATERIALS AND METHODS

### Library preparation and sequencing

Capture-based enrichment, followed by sequencing, was used to evaluate the impact of input material parameters on data quality metrics and the performance of biomarker detection. For control of hybrid capture robustness, 50 ng DNA of a reference cell line (NA24149, Coriell Institute) was converted into a sequencing library using NEBNext Ultra II DNA Library Prep (New England Biolabs). 200 ng of libraries were used for hybridization capture. 114 FFPE tissue samples derived from patient solid tumors previously processed with an amplicon-based assay (ArcherDx Lung, IDT) were utilized to prepare sequencing libraries. A pathologist reviewed all FFPE samples to estimate tumor cell content. RNA was extracted from FFPE scrolls using Reliaprep FFPE Total RNA Miniprep (Promega). Another preparation was conducted for a subset of samples, omitting DNase addition and yielding total nucleic acid (tNA). A portion of each tNA sample was further purified using RNA Clean & Concentrator (Zymo Research) to obtain matched RNA samples. Simultaneously, another portion of tNA sample underwent DNA isolation with genomic DNA Clean & Concentrator (Zymo Research). RNA isolated from FFPE-treated reference cell lines (GM24143, GM24149, GN24385, Coriell and Seraseq FFPE Tumor Fusion RNA v4 Reference Material, Seracare) were used as controls. DV200 was determined as the RNA integrity measure.

For the majority of analyses, 50 ng of RNA input was used. For a subset of samples, lower input amounts (10 and 20 ng) were used to assess the impact of input quantity on the quality metrics. Moreover, two samples were tested in 4 replicates at 2, 5, 10 and 20 ng inputs to test reproducibility at low inputs. For the amplicon assay (ArcherDx Lung, IDT) head-to-head comparisons with RNAtarget Oncology Solution, 50 ng (clinical samples) and 20 ng (reference samples) of RNA were used as input, processed according to manufacturer recommendations. Following the manufacturer’s instructions, the TruSight Tumor 170 (TST170, Illumina) assay was executed on 50 ng of RNA.

RNA or tNA were processed using RNAtarget Oncology Solution (ROS, SOPHiA GENETICS) at amounts ranging from 2 to 50 ng (quantity indicated for each experiment). Briefly, the input material was reverse transcribed using a mixture of random hexamers and 16 bp long polyA oligonucleotides. Sequencing adapters were ligated, and the libraries were amplified with PCR. Sequencing libraries were prepared from 50 ng of DNA isolated from various reference cell lines (Coriell) using Solid Tumor Solution (SOPHiA GENETICS) for capture efficiency tests. Target enrichment was performed by an overnight hybridization capture at 65°C with ROS panel. Post-capture libraries were amplified using 15 PCR cycles and sequenced on NextSeq 550 sequencer (Illumina) aiming to achieve an average depth of 1.5M read pairs per sample.

### Detection of fusions, exon skipping, short variants and expression measurement

Sequencing data were processed with an analytical pipeline of SOPHiA GENETICS for RNAtarget Oncology Solution. Reads were aligned to genome assembly hg19 (GRCh38.p13) with STAR aligner (STAR-2.7.0f)^27^. All coverage calculations considered molecules rather than reads, where a single molecule is defined by unique mapping start and end position, regardless of the number of sequenced fragments with the exact coordinates. Fusions were detected using a Bayesian caller. In cases where several aberrant transcripts were detected for an event (fusion or exon skipping) that involved the same genes and exons, the most likely event was chosen based on the highest molecular support. The fraction of aberrant transcript molecules for each event was calculated relative to all captured molecules at the junction. Reciprocal fusions were excluded from the analyses. The limit of detection was defined as the number of event-supporting molecules present in the input required for detection of at least 10 molecules with 95% sensitivity. To account for possible variability, the upper confidence interval (95% quantile) was calculated and considered as the limit of detection value.

Small variants were detected based on the molecular support and subsequently filtered based on the variant allele fraction (VAF, minimum 3%), minimal read and molecule support, background noise, displayed strand bias, presence outside of the target regions or in regions known to be problematic or displaying significant noise.

Gene expression was evaluated by calculating median molecular coverage of targeted exon-exon junctions normalized by the geometric mean of the molecular coverage obtained across targeted exon-exon junctions of six control genes.

### Fusion fragment mapping evaluation

Artificial reference sequences representing the chimeric transcripts identified in each of the analyzed samples, were generated in-silico. These served to align the sequencing reads of each corresponding sample using bwa paired end^28^. Similarly, control sequences representing the non-chimeric transcripts were generated and used to align the same sequencing datasets. The alignment aimed to map fragments over a window of 1000 basepairs around the breakpoint (for the chimeric transcript) or up to the ends of the exons adjacent to the position corresponding to the breakpoint (for the control transcript). Using the mapping coordinates of the fragments over the artificial reference sequences, fragments overlapping the breakpoint were identified and their size and distance to the breakpoint size were measured.

### Quality metrics calculation

Control gene coverage was determined as the median count of molecules detected across targeted NRF1 regions. The on-target rate reflects the fraction of base pairs falling within the panel target region relative to all base pairs in mapped reads. Read group size was calculated by aggregating all sequenced fragments by their mapping start and end position and counting duplicates. DNA content metric was evaluated by comparing median molecular coverage over representative exon-intron junctions of the control gene.

## RESULTS

### The robustness of RNA capture panel can be estimated using DNA input

To assess the impact of various technical parameters on the results reported from targeted RNA sequencing, we used RNAtarget Oncology Solution (ROS), a hybridization capture-based NGS test that targets 45 genes frequently mutated in lung cancer and other solid tumors (**Supplemental Table 1**) and allows for detection of fusions, exon skipping events and small variants as well as gene expression measurement (**Figure 1A**).

**Figure 1.**
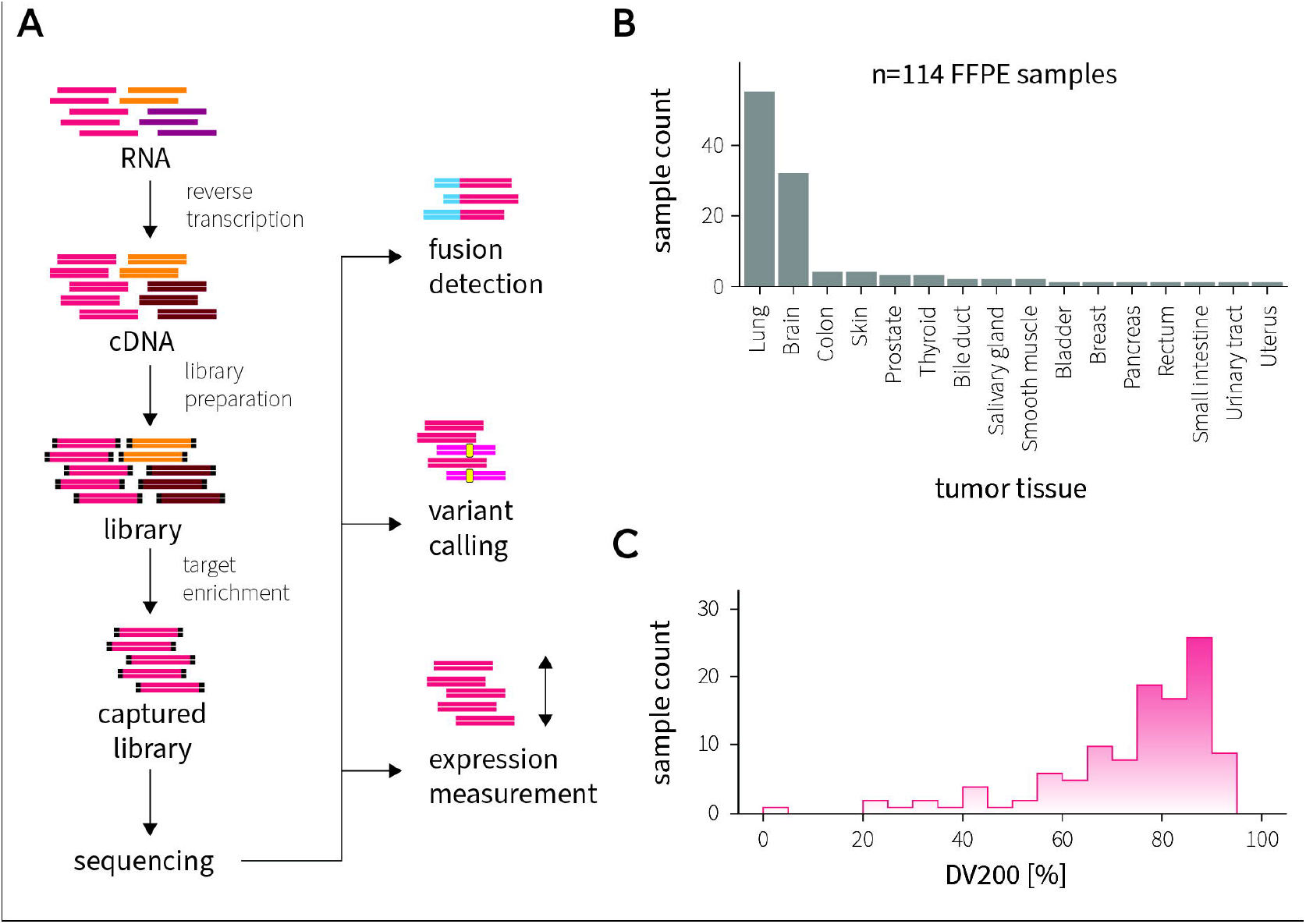
Overview of RNA processing workflow. **(A)** Schematic representation of targeted RNA-seq data generation using capture-based enrichment. **(B)** Composition of the cohort of clinical FFPE samples used in the analyses. **(C)** Overview of sample fragmentation evaluated by DV200 measurement.

Achieving effective hybridization capture performance relies on robust pull-down of all the regions targeted by the panel. However, assessing robustness can be challenging with RNA as the targets are not present at equimolar amounts. Consequently, evaluation of target capture performance was executed on libraries prepared from DNA of a reference cell line. Despite the imperfect probe overlap with a subset of targets due to presence of noncoding regions in DNA, the obtained coverage profile displayed good overall uniformity (**Figure S1A**). Coverage dropout was observed only for several DNA regions with less than 60 bp overlap with the probes (**Figure S1B**) and did not depend on the GC content (**Figure S1C**). In summary, DNA library capture proves to be a robust method for testing capacity for uniform target enrichment of RNA-targeting panels.

### Molecular coverage of the control gene is a good proxy of input material conversion

ROS assay was used on material isolated from a cohort of clinical specimens consisting of FFPE solid tumor tissue. The tested cohort comprised of 114 RNA samples from 16 different tissues (**Figure 1B**). Samples displayed various extent of fragmentation, with approximately half of them having DV200 below 80% (**Figure 1C**).

For constant input quantities, the obtained library yield should give an approximate measure of the conversion rate of RNA molecules to sequencing libraries, which in turn depends on RNA quality. However, the correlation between DV200 and the obtained sequencing library concentration was only moderate (Pearson R^2^=0.25, p<10^-6^, **Figure S2A**) suggesting that fragmentation alone is not fully predictive of the conversion. Conversely, the median molecular coverage of a control housekeeping gene, NRF1, correlated well with the library concentration (Pearson R^2^=0.42, p<10^-14^, **Figure 2A**) showing that it can be used as a good proxy of the conversion rate derived directly from the sequencing data. NRF1 was chosen due to its stable expression across a wide range of tissues (www.gtexportal.org/home/). In addition, molecule count of the control gene scaled with the input amount (**Figure S2B**) and was consistent across replicates (**Figure S2C**) as observed in data acquired from varying RNA quantities. An external laboratory processed a subset of samples to test the control gene coverage reproducibility further. The results demonstrated strong correlation of control gene coverage values measured at both sites (Pearson R^2^=0.91, p <10^-7^) (**Figure S2D**). Therefore, we concluded that NRF1 molecule count is a robust of RNA conversion measure.

**Figure 2.**
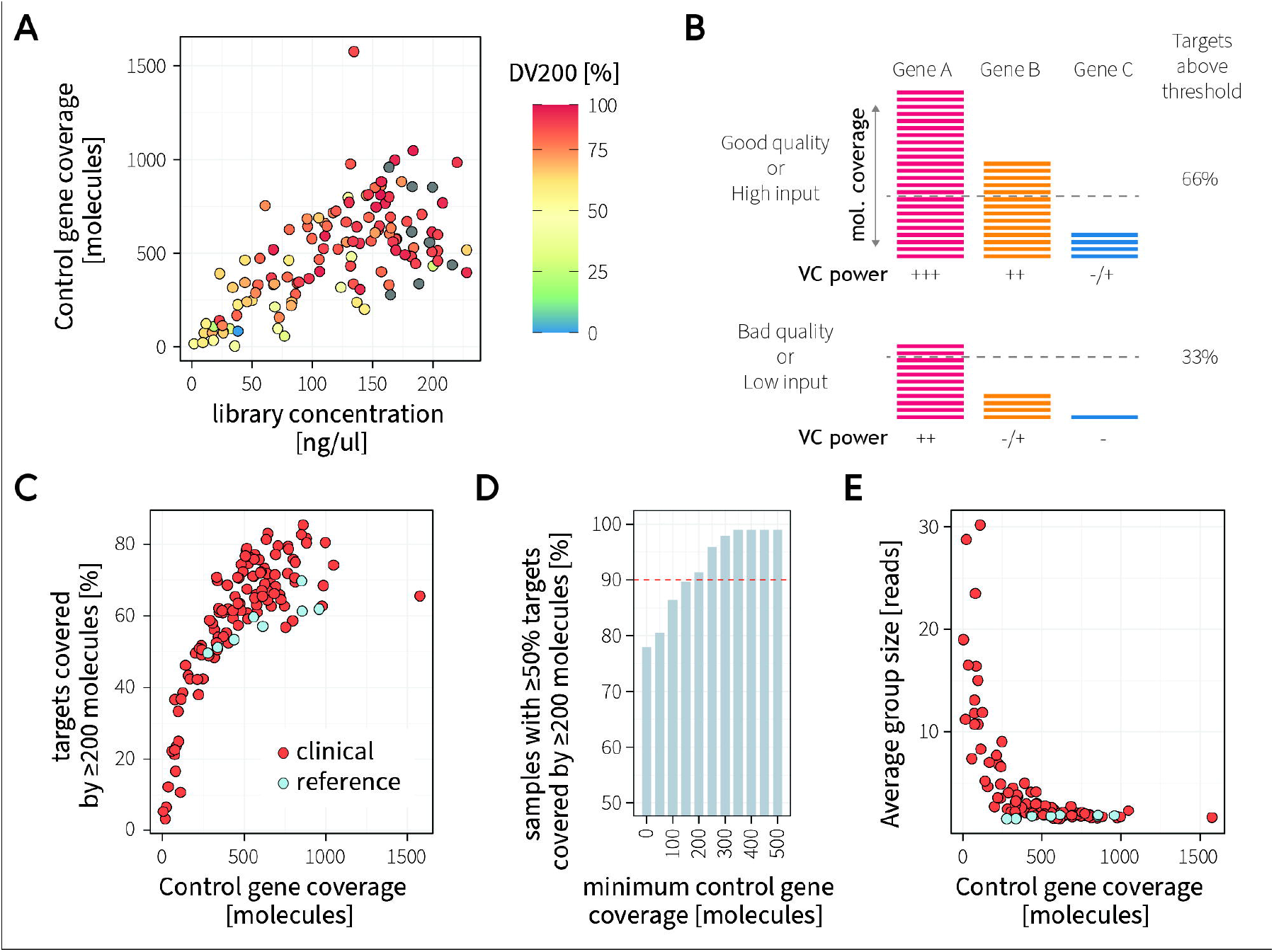
Control gene signal is a good proxy of data quality. **(A)** Control gene coverage (y axis) as a function of library concentration (x axis) obtained in the tested samples; color reflects the measured DV200 value. **(B)** Schematic representation of the impact of sample quality and quantity on the fraction of well covered regions. **(C)** Fraction of targets covered by at least 200 molecules (y axis) as a function of control gene coverage (x axis) in all tested samples. **(D)** Fraction of samples that had at least half of their targets covered by at least 200 molecules depending on the threshold of control gene coverage for sample inclusion. **(E)** Average read group size (y axis) as a function of the control gene coverage (x axis) in all tested samples.

### Control gene coverage correlates with multiple data quality features

At least 75% of the sequenced basepairs fell onto the target regions in 87% (99/114) of the samples (**Figure S2E**) despite the absence of ribosomal RNA (rRNA) depletion step during sample processing. Samples with lower on-target rates displayed lower control gene coverage (CGC) (**Figure S2F**), which suggested that the overall material available to be captured by the panel is linked to molecular diversity of the sample.

Good sensitivity of detection of genomic alterations requires sufficient coverage of individual targets. For example, detecting small variants at an allelic frequency of 5% with 95% sensitivity requires coverage of at least 200 molecules, assuming that at least 5 sequenced molecules should support variant presence. Since different transcripts are present at varying amounts in a given quantity of RNA, it is crucial to control the coverage of individual targets (**Figure 2B**). The fraction of sufficiently covered targets for the tested panel was closely correlated to the CGC (**Figure 2C**). Ensuring that at least half of the targets are covered by 200 molecules in 90% of the samples required CGC of at least 150 molecules (**Figure 2D**). As expected, the fraction of well-covered regions for a given sample scaled with the input material quantity (**Figures S2G** and **S2H**).

Measuring PCR duplicates (read group size) is an important indicator of whether a library may benefit from deeper sequencing. If the mean group size significantly exceeds one read, one can assume that all input molecules were sequenced and a further increase in sequencing depth will not significantly improve the performance. We observed that samples with low CGC displayed high average group size and vice versa (**Figure 2E**). This observation confirms that CGC is a quality metric that reflects the molecular diversity of the samples.

In summary, the molecular coverage of the selected control gene correlates with several key metrics (input material fragmentation, on-target rate, target coverage, molecular diversity) and therefore can serve as a good proxy of overall data quality.

### Reliable expression measurement requires a DNA-aware approach

To facilitate expression comparison across samples, median molecule counts calculated per gene were normalized to those measured for control genes (see **Materials and Methods**). Such normalization allows to obtain consistent expression measure independent of the molecular diversity of the sample. Indeed, we observed that normalized expression, contrary to molecule counts, was not correlated with CGC (**Figure 3A**) and did not depend on the input quantity (**Figure S3A**).

**Figure 3.**
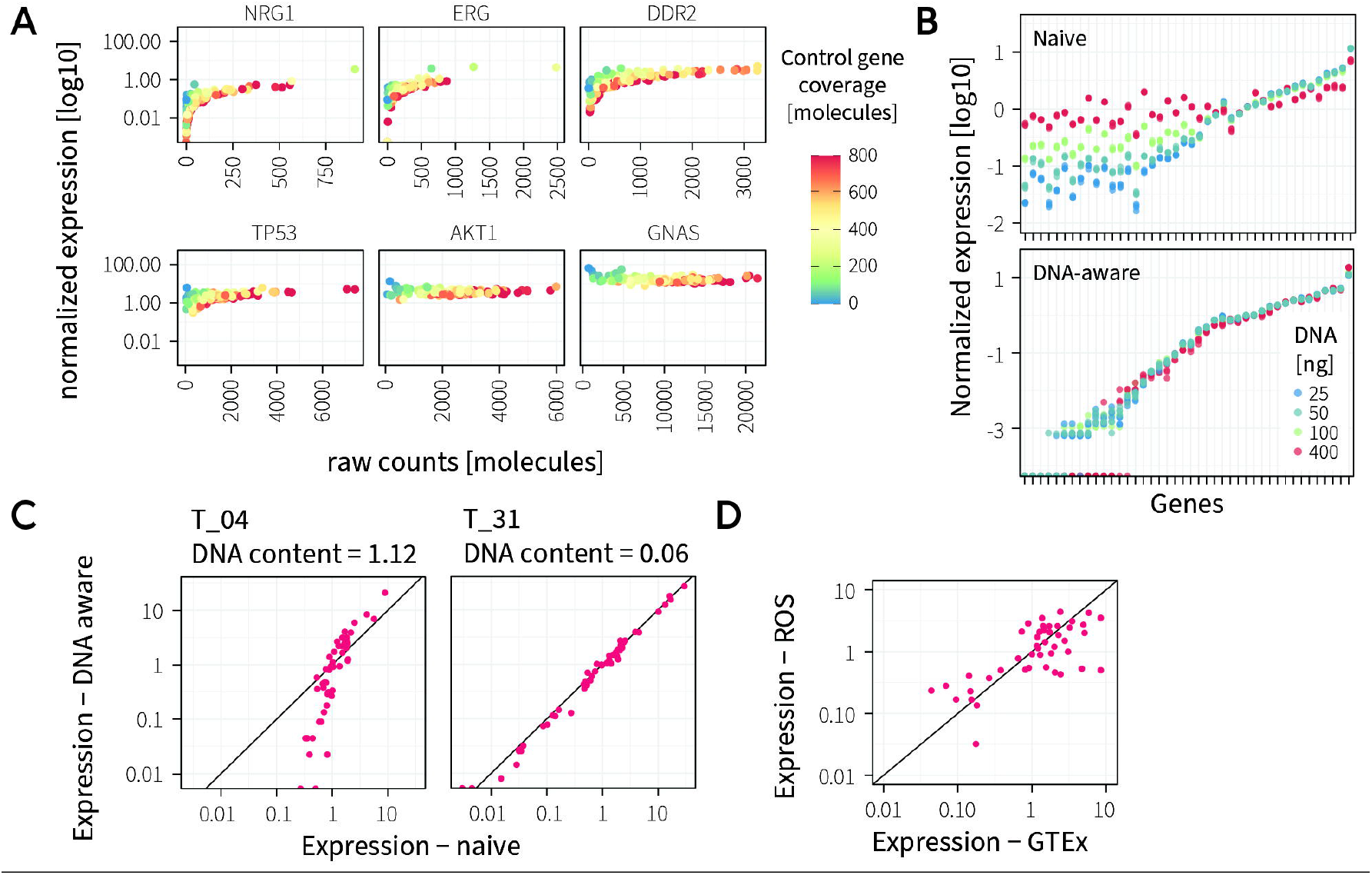
Accurate expression measurement requires DNA-aware estimation. **(A)** Gene expression signal across all tested samples shown as normalized (y axis) versus non-normalized (x axis) molecule counts for 6 representative genes; color indicates control gene coverage in each sample. **(B)** Normalized gene expression values measured in a reference RNA sample containing varying DNA quantities with a naïve (top) or DNA-aware (bottom) method. **(C)** Normalized expression measured in two clinical samples containing varying DNA quantities with a naïve (x axis) or DNA-aware (y axis) method. **(D)** Median normalized gene expression values measured with ROS in lung tumor samples (y axis) and reported in the GTEx database (x axis).

DNA is more stable than RNA and is a frequent contaminant of RNA preparations. To investigate the impact of DNA on measurement of gene expression levels we processed RNA of a reference sample with varying amounts of DNA admixture. We observed that expression measured as median-normalized count of all unique molecules presented less variation in the presence of a larger quantity of DNA (**Figure 3B, top**). In the presence of DNA, values for highly expressed genes were underestimated and the opposite was observed for poorly expressed genes and the extent of this bias was proportional to DNA quantity (**Figure 3B, top**). To account for this effect, we applied a measurement that does not consider presence of reads potentially derived from DNA. Application of DNA-aware molecule counts yielded reproducible results regardless of DNA quantity in the samples (**Figure 3B, bottom**).

We developed a DNA content calculation based on the molecular coverage ratio within 10 bp of the exon-intron junctions to inform the sample of excess DNA present. We compared expression measured with “naïve” and DNA-aware methods in clinical samples from which both RNA and a mixture of RNA and DNA (total nucleic acid, tNA) were isolated. The results revealed significant differences in the estimated expression for a sample with a high DNA content metric. Conversely, little or no effect was observed in sample containing low DNA quantity (**Figure 3C**).

We then applied DNA-aware expression measurement across all processed samples. We observed clustering according to the source tissue based on the normalized expression values (**Figure S3B and S3C**). Large differences were observed between expression in the lung and other tissues for several genes, such as ROS1 and NTRK2 (**Figure S3D**). When median expression in the lung samples was compared to that extracted from the GTEx database (www.gtexportal.org/home/), highly significant correlation was observed (Pearson R^2^=0.7, p<10^-13^), despite comparing values from tumor samples to those derived from healthy lung tissue (**Figure 3D**).

In summary, these observations suggest that DNA presence may severely impact gene expression measurement. It is important to estimate the sample DNA content and use DNA-aware expression measurement for reliable results.

### Single fusion partner targeting allows to efficiently detect diverse chimeric transcripts

Data generated with ROS on clinical and reference samples was used to detect gene fusions and exon-skipping events. The results were compared to those previously obtained using a nested amplicon-based approach (Archer FUSIONPlex). In case of discordant results, a third assay, TST170 (Illumina), was used to determine the ground truth (**Supplemental Methods and Supplemental Table 2**). ROS assay demonstrated 100% sensitivity (84/84 events, including 70/70 in 66 clinical samples), detecting of 7 events missed by the amplicon but confirmed by TST170 (**Figure S4**). Events detected at molecule counts below 10, used as a threshold for confident calls, invariably represented a small fraction of total molecules at the fusion junction (**Figure 4A**) suggesting artefactual origin. True positive calls displayed a broad distribution of the fractional values. A low number of molecules supported two events not confirmed by TST170.

**Figure 4.**
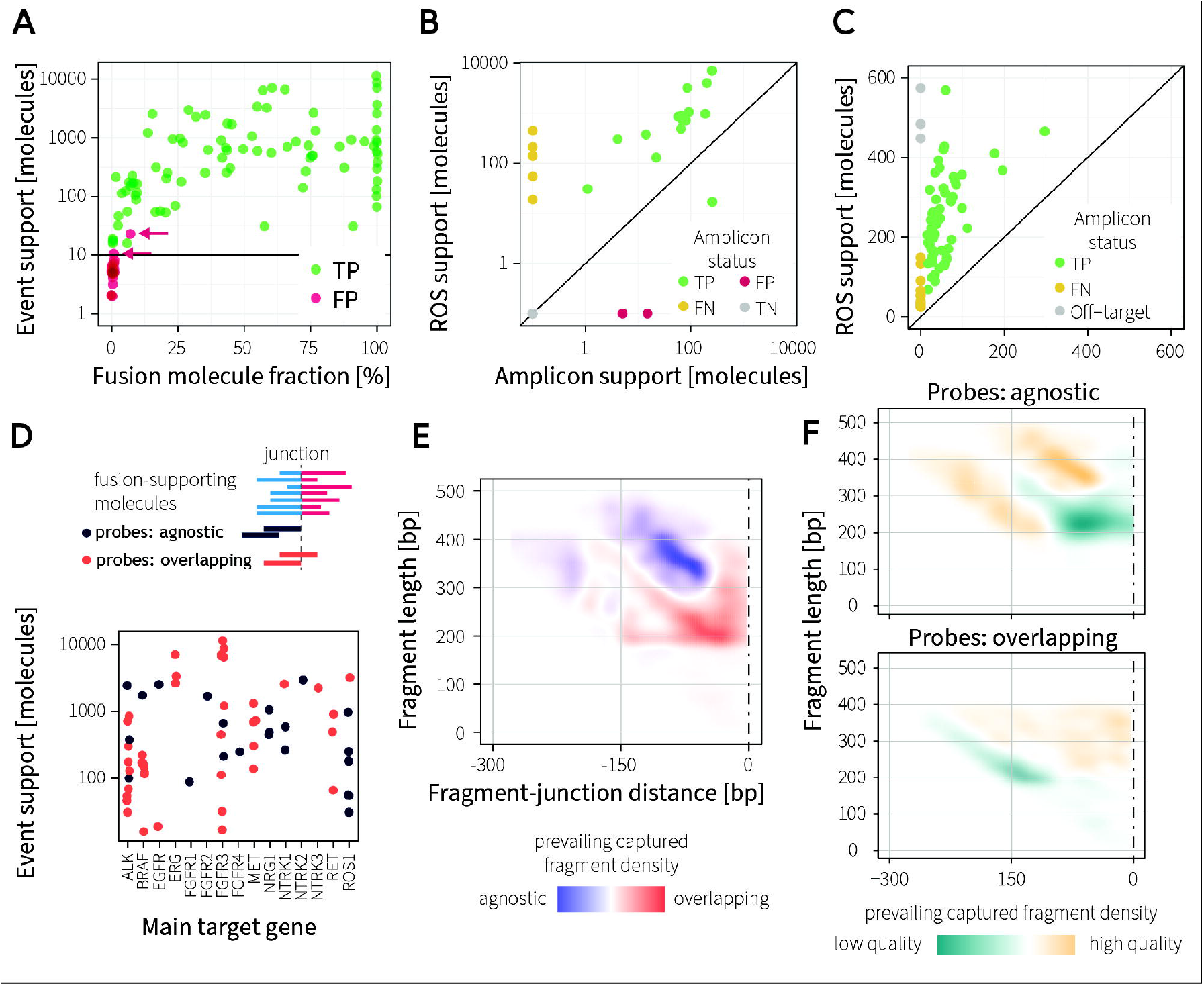
Efficient aberrant transcript detection with single partner targeting. **(A)** Molecule counts as a function of fraction of all molecules found at the breakpoint location for all identified fusion and exon skipping events; threshold for high confidence detection is set at 10 molecules; red arrows indicate false-positive calls. **(B)** Supporting molecule counts for identified events split by the main target gene; color indicates the fusion capture approach. **(C)** Molecule counts for events identified in 23 clinical samples (50 ng RNA) with amplicon assay (x axis) and ROS (y axis); color indicates call status in the amplicon data: TP true-positive, FP false-positive, FN false-negative, TN true-negative. **(D)** Molecule counts for events identified in reference samples (20 ng DNA) with amplicon assay (x axis) and ROS (y axis); color indicates call status in the amplicon data. **(E)** Heatmap representing difference in the density of fragments that span the fusion transcript junction, depending on the capture approach; fragments are displayed according to their position (X axis) and size (Y axis); blue indicates prevalence in agnostic, and red in overlapping capture. **(F)** Heatmap representing density of fragments depending on sample quality, stratified by control gene coverage below (low quality) or above (high quality) the median value, for agnostic (top) and overlapping (bottom) targeting.

To investigate the discrepancies between capture and amplicon results, for a subset of samples (n=23) where sufficient material was available, the amplicon test was repeated and the number of unique molecules (i.e. amplicons) supporting the events was compared to the one obtained with ROS (**Figure 4B**). The molecule count was higher with ROS for 19/20 true events. Five of the tested events missed by the amplicon were also not detected during the repeated test. To expand the comparison, we processed 3 types of reference samples, each bearing multiple fusion or exon skipping events, in replicates, using both amplicon and ROS assays. Consistently, higher molecule counts were observed for ROS across all detected events (**Figure 4C**). These observations suggest that converting input RNA to sequencing libraries was more efficient for capture-based assay than amplicon.

ROS targets chimeric transcripts in a partner-agnostic manner but for selected frequent fusions, also the partner exons are targeted to potentially improve the detection sensitivity. Molecule counts for fusions detected in a partner-agnostic manner were similar to those where both partners were targeted (Wilcoxon test p-value=0.89) (**Figure 4D**), suggesting comparable capacity to capture chimeric transcripts. Despite the efficiency of partner-agnostic capture, probe positioning in this approach resulted in missing fragments starting close to the chimeric transcript breakpoint, favoring long fragments (>300 bp-long) with at least ∼75 bp overlap with the captured fusion partner (Figure 4E, blue density). In comparison, junction-overlapping targeting allowed for the enrichment of shorter fragments, starting closer to the junction (**Figure 4E**, red density). Moreover, low sample quality was related to decreased capture efficiency of specific of molecules. For agnostic targeting, this was pronounced for fragments <300 bp (**Figure 4F**, top) while for junction-overlapping targeting – for fragments located further away (>120 bp) from the junction (**Figure 4F**, bottom). These results suggest that higher material quality provides overall better sensitivity for agnostic capture by efficient recovery of smaller fragments at the fusion proximity. In contrast, for overlapping capture the benefit is restricted to fragments having an overall small overlap beyond the breakpoint.

### Factors affecting the limit of detection of aberrant transcripts

For a given quantity of RNA, the number of detected molecules is determined by the sample conversion rate and abundance of the particular transcript. The latter depends on the source tissue and tumor content. Indeed, we observed that the number of detected molecules, for most events scaled with the control gene molecule count (**Figure 5A**), tumor content (**Figure S5A**) or the input material (**Figure 5B**).

**Figure 5.**
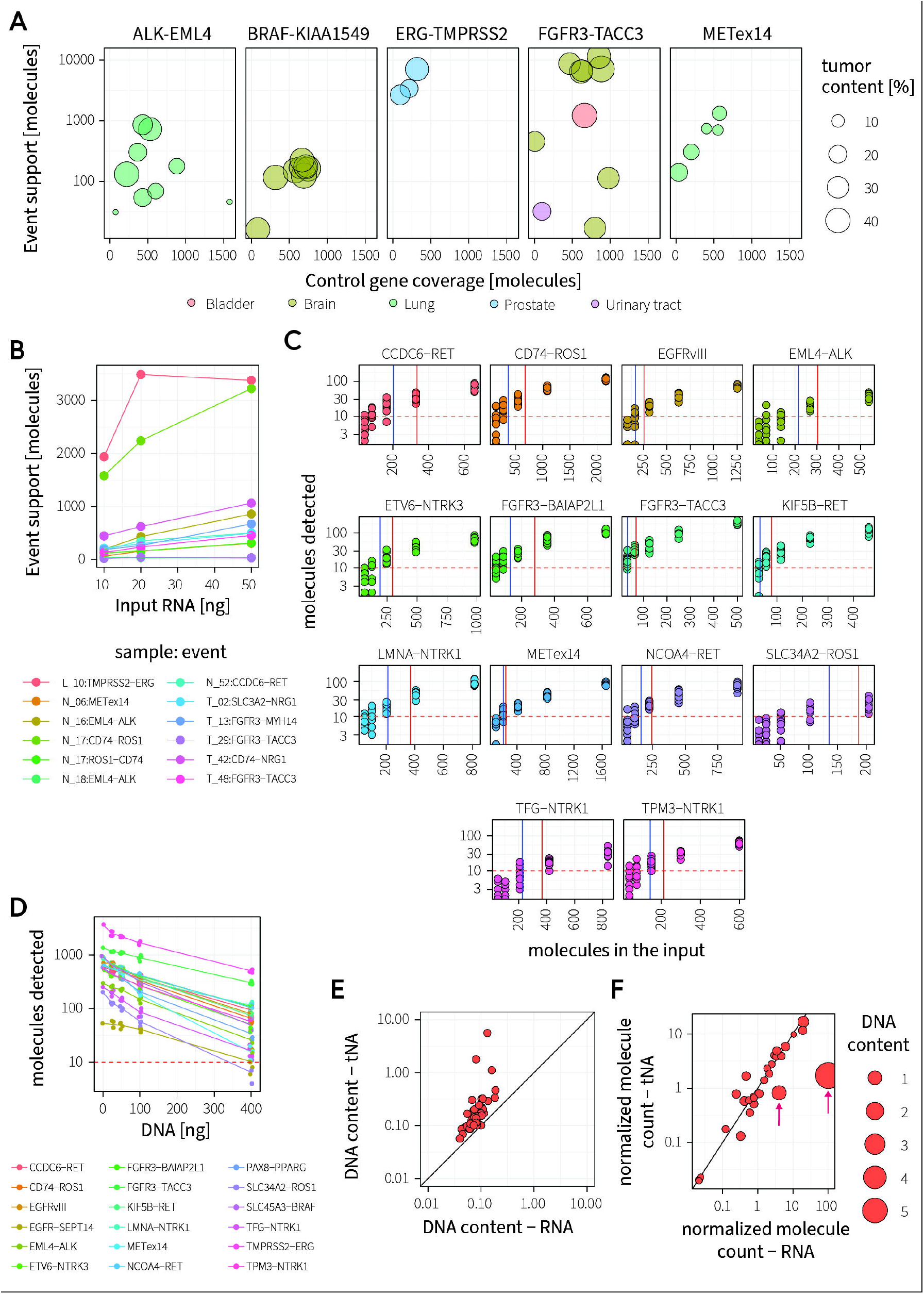
Key factors affecting aberrant transcript detection. **(A)** Supporting molecule count (y axis) as a function of control gene coverage (x axis) for events present in at least three clinical samples; points are colored according to source tissue; point size reflects tumor content. **(B)** Supporting molecule count (y axis) as a function of input RNA (x axis) for events present in clinical samples that were processed with varying input amounts. **(C)** Limit of detection estimation displayed as detected supporting molecule counts (y axis) versus expected molecule counts in the input material (x axis) measured in replicates of a serial dilution of a reference sample for 14 distinct targeted events; blue vertical line indicates quantity at which 95% sensitivity was expected according to probit analysis; red vertical line indicates the 95% confidence interval for the sensitivity, established as the limit of detection. **(D)** Supporting molecule count (y axis) as a function of DNA (x axis) added to 50 ng of a reference RNA sample. **(E)** Correlation of estimated DNA content for pairs of preparations of either RNA (x axis) or tNA (y axis) extracted from the same sample. **(F)** Control gene coverage-normalized molecule count for events detected in RNA (X axis) and tNA (Y axis); size of points reflects DNA quantity; arrows indicate events found in samples with highest DNA content.

To investigate the limit of detection (LoD) of the assay, a known number of targeted molecules should be used as input. We performed a test where a reference sample carrying 14 distinct targeted events, with quantified molecule counts provided by the manufacturer, was serially diluted in background RNA. These dilutions were used in replicates as input for ROS. A rigorous statistical test was used to evaluate the LoD (see **Materials and Methods**). The calculated values varied across different events but values for 12/15 of them were within a relatively narrow range of 186-370 molecules (**Figure 5C, Supplemental Table 3**). To validate the LoD range, we repeated the experiment with serial dilutions of a clinical sample bearing CD74-NRG1 fusion, targeted in a partner-agnostic manner, with the number of fusion molecules per ng of RNA estimated using ddPCR. The LoD for this event was 357 molecules, indicating that estimates from the reference sample can be translated to events present in clinical samples.

The effect of DNA presence on the detection of pathological transcripts was investigated in an experiment where a range of DNA quantities were present in a processed reference sample. We observed that high DNA content impacts the sensitivity of the chimeric transcript detection by decreasing the number of detected supporting molecules (**Figure 5D**). The DNA content metric was higher in tNA isolated from the clinical samples than in corresponding RNA samples (**Figure 5E**), which validated this quality indicator. For samples with little DNA, the ratio between molecule counts supporting detected events in the tNA versus RNA sample was proportional to such ratio calculated for the control gene (**Figure 5F**). However, the relative decrease in fusion-supporting molecules in tNA was much stronger in the two samples with the highest measured DNA content.

In summary, our results highlight the importance of DNA content as an essential quality metric. Results from samples with high DNA content should be evaluated carefully due to the risk of false-negative results of chimeric transcript detection.

### The reliability of short variant calling from RNA depends on target expression

The capability of short variant detection from RNA was evaluated on a subset of the FFPE specimen (n=57) for which corresponding DNA was available for testing. To establish the ground truth, DNA was processed with a capture-based assay targeting regions partially overlapping with the targets of ROS. Samples of fully characterized cell lines from the Genome in a Bottle consortium were processed as controls. Variants detected at an allele frequency of 5% or above in the DNA analysis were monitored in the RNA-derived data.

The correlation of VAF between values measured in DNA and RNA was highly significant (Pearson R^2^=0.82, p<10^-15^) (**Figure 6A**). Some variants demonstrated significant difference in VAF, possibly due to allelic imbalance. For targets covered by at least 200 molecules in the RNA dataset, 99.7% of variants (312/313) were detected, including 99.7% (300/301) in the clinical samples (**Figure 6B**). When the analysis was expanded to all variants present in the common regions, regardless of coverage, the sensitivity was 91.4% (491/537) for all samples and 93.5% (458/490) for the clinical samples. Additional variants were detected in RNA, but the vast majority (119/143, 83.2%) either displayed VAF below 5% or was present in regions with coverage below 200 molecules (**Figure 6B** and **6C**).

**Figure 6.**
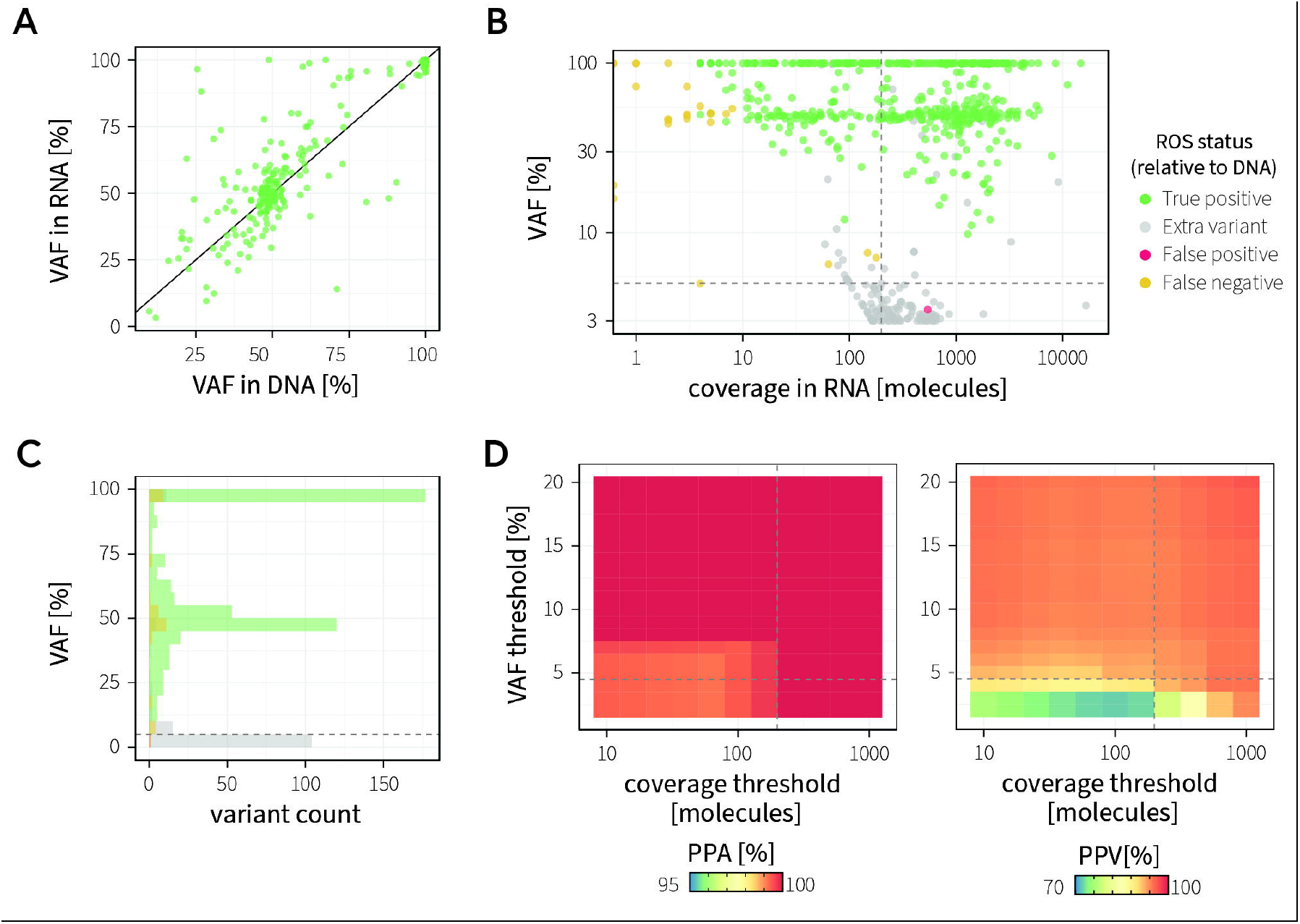
Variant calling performance in RNA is defined by coverage and VAF. **(A)** Correlation of VAF measured in DNA (X axis) and RNA (Y axis). **(B)** All detected variants depending on the molecular coverage (X axis) and VAF measured in RNA (Y axis). **(C)** Distribution of variant counts depending on their VAF measured in DNA for true positive calls (green), false negatives (red) and extra variants (grey). **(D)** Heatmaps showing PPA (left) and PPV (right) values depending on the threshold used for variant calling on minimal coverage (X axis) and VAF (Y axis).

Variants not detected by ROS were characterized by low coverage in RNA and/or low VAF in DNA (**Figure 6B** and **6C**). Conversely, many alterations with VAF>10% were successfully detected even in targets with coverage below 200 molecules. Both positive predictive agreement (PPA) and positive predictive value (PPV) varied depending on the coverage at different VAF thresholds (**Figure 6D**). This was a direct consequence of the fact that (i) at small VAF values, lower coverage is sufficient to detect alterations and (ii) extra variants were present mostly at low fractions. Moreover, variant detection sensitivity was linked to the fraction of target regions covered by at least 200 molecules as demonstrated by comparison of numbers of true-positive calls detected in the same samples with 10, 20 and 50 ng RNA input (**Figure S6**). In summary, these observations underscore the strict dependency of variant calling on molecular coverage of the target in RNA-seq data.

### DNA presence may be beneficial for short variant detection

We tested how the presence of DNA impacts variant detection by analyzing results from the tNA samples. We observed that molecular coverage of variant positions was higher in tNA than in RNA (**Figures 7A and 7B**). In the experiment with DNA admixture to the reference sample, we observed that increasing amounts of DNA added to the reference sample positively impacted the fraction of well covered regions (**Figure S7A**). Consistently, 14 additional variants, confirmed by DNA-only analyses, were detected in tNA owing to higher coverage (**Figure 7A**). The correlation of tNA VAF values with those measured in DNA was marginally higher (Pearson R^2^=0.87, p<10-^15^) compared to the correlation measured for RNA (R^2^=0.82), suggesting a predominant role of RNA molecules in support of detected variants (**Figure 7C**).

**Figure 7.**
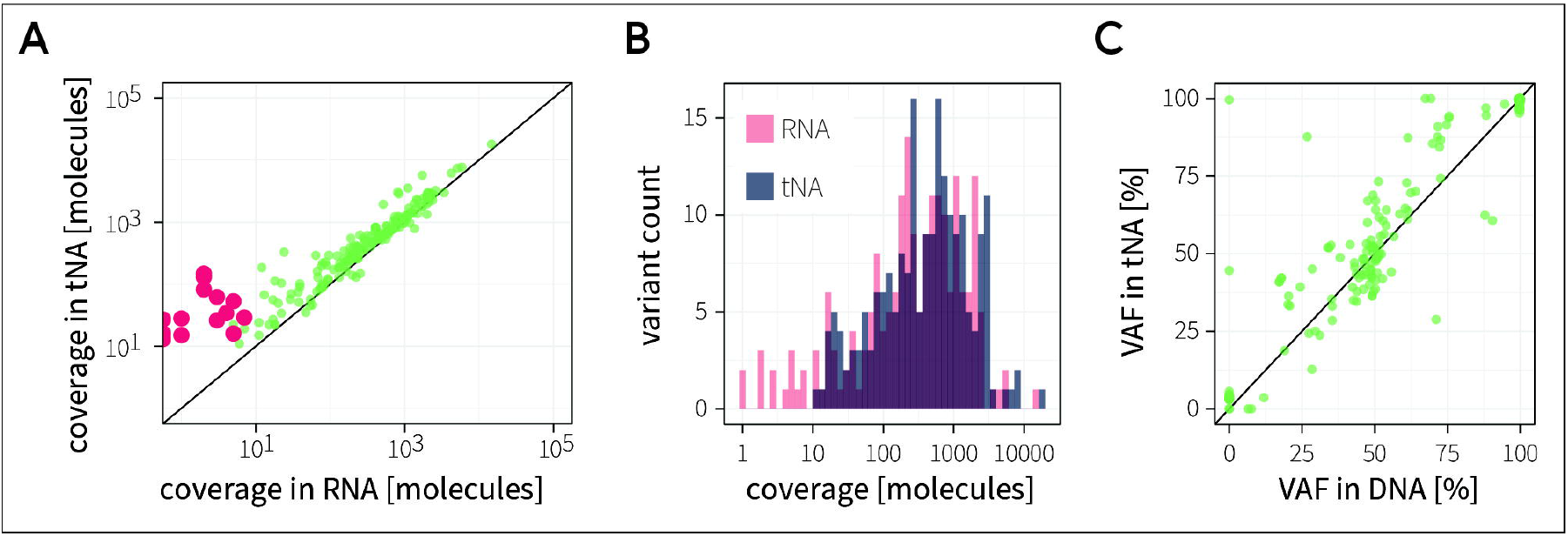
DNA presence allows to detect additional variants comparing to RNA. **(A)** Coverage comparison at variant positions in RNA (X axis) and tNA (Y axis); variants detected in tNA only displayed in red. **(B)** Distribution of coverage at variant positions for tNA and RNA datasets. **(C)** Correlation of VAF measured in DNA (X axis) and tNA (Y axis); false-negative variants displayed in red.

These results indicate that DNA presence may be beneficial for the sensitivity of variant detection, particularly in lowly expressed target genes. However, as shown above, large amounts of DNA in the sample may hamper the detection of aberrant transcripts; therefore, DNA content should be closely monitored.

## DISCUSSION

### Control of molecular diversity as a universal quality measure

We present a thorough investigation of parameters that affect the efficiency of concurrent detection of several types of clinically and biologically relevant information in targeted RNA-seq data. Previous studies put a lot of emphasis on the input material features and were typically conducted from the perspective of a single type of biomarker: gene expression^17,29–33^, fusions^34,35^ or short variants^36,37^. We describe universal data-based quantitative measures that aid in correct interpretation of multiple types of results (**Table 1**).

**Table 1.** Critical parameters for the control of performance of detection of various biomarker types in targeted RNA sequencing data.

|  | Parameter | Recommendations and remarks |
| --- | --- | --- |
| General | RNA conversion rate | maximize, evaluate by control gene coverage |
|  | On-target rate | maximize |
|  | Group size | recommended ~2-5 reads, evaluate by control gene coverage |
|  | DNA content | control with a dedicated metric |
| Expression | Normalization | use housekeeping genes for normalization |
|  | DNA content | use DNA-aware expression measure |
| Aberrant transcripts (fusions, exon skipping) | Detection threshold | 10 molecules for ROS, should be evaluated based on noise levels |
|  | DNA content | higher DNA content has negative impact on detection sensitivity |
| Variant detection | Fraction of covered targets | maximize, evaluate by control gene coverage |
| | Target coverage | depending on VAF, e.g. $\geq 200$ molecules for VAF=0.5%, $\geq 100$ molecules at VAF=10% |
|  | DNA content | higher DNA content has positive impact on detection sensitivity |

The detection chance of a rare event depends on the sum of observations. Therefore, the number of interrogated RNA molecules should be maximized to ensure good sensitivity of detecting a genomic alteration by RNA-seq. Undoubtedly, the major obstacles to reaching high molecular diversity are the RNA quality, quantity, and conversion rate achieved with the applied assay^38^. RNA integrity, expressed in the DV200 value, was shown to correlate with sequencing library output better than alternative metrics^39^. Nevertheless, we observed that fragmentation did not serve as an optimal proxy of the RNA conversion or the overall library diversity. Conversely, a single metric, the molecule coverage of a control housekeeping gene, correlated with various quality measures: the library yield, on-target rate, duplicate rate, fraction of well-covered targets, number of alteration-supporting molecules and variability of expression measure.

We observed that poor quality samples displayed large read group sizes, reflecting high PCR duplicate rates. This indicates that such samples would not benefit from additional sequencing, as little or no new molecules could be potentially identified, which is consistent with previous observations that deeper sequencing does not improve fusion detection sensitivity^40^. In practical terms, using lower quality material is similar to using lower quantities of RNA. We show that molecule counts, and consequently the performance, scale with both features and the increase in the number of molecules can be achieved by augmenting the input quantity.

In addition to the input material parameters, the efficiency of RNA conversion is also influenced by the employed library preparation workflow^41^. We observed good on-target rates across FFPE samples despite the lack of rRNA depletion step in the tested workflow, suggesting that target enrichment acts as the rRNA depletion step. Moreover, hybridization-based enrichment reproducibly reported higher molecule counts than a nested amplicon-based assay for fusions and exon skipping events in both clinical and reference samples, consistently with a previous report on fusion detection sensitivity^42^.

### Impact of DNA presence on targeted RNA-seq results

We demonstrated that the presence of DNA significantly impacts the detection of various biomarkers, thus is a parameter important to control. The undesired effects on the result interpretation included biasing expression values and decreasing the number of detected aberrant transcript molecules. At the same time, DNA presence can improve detection of variants in poorly expressed genes. We developed a DNA content metric whose value correlates with several output features, such as the expression bias and the number of alteration-supporting molecules. Its application allows to better interpret the results by informing about the tested sample composition and the potential impact.

The effect of DNA contamination on gene expression measurement can be counteracted by applying a method that can unambiguously identify the RNA-derived reads. We demonstrated that values obtained with DNA-aware measurement are insensitive to DNA presence in the sample.

### Insights for detection of different biomarker types

Previous studies reported limited correlation of measured expression between replicates or matched fresh-frozen and FFPE samples in RNA-seq^40,43^. Conversely, we demonstrated a significant correlation between values measured for a small subset of genes in FFPE tumor material to those reported for lung tissue measured in fresh-frozen samples. We conclude that reliable expression estimation requires efficient normalization and an approach sensitive to DNA presence in the sample. Further improvement may be possible by applying molecular barcoding to resolve fragment collisions for molecule calculations of the abundant transcripts^38,44,45^.

The efficiency of detection of aberrant chimeric transcripts is known to vary across targets^46^. We demonstrated that partner-agnostic fusion detection with hybridization capture has comparable efficiency to an approach in which both partners are targeted. The number of detected molecules supporting each event was related to the tumor content and the input conversion rate and was higher than measured for a nested amplicon method. Interestingly, a study utilizing an the nested amplicon assay reported LoD values lower than the ones we estimated for hybridization capture^34^. There were several differences in the approach: the method applied for input molecule number determination was different, we used technical replicates and applied a rigorous statistical test, which makes the LoD value conservative. Nevertheless, the LoD as a number of molecules is of little practical use as the expected value in the starting material cannot be readily determined and LoD does not correlate with fusion detection sensitivity^47^.

It has long been recognized that RNA-seq allows to identify small variants in cancer and hereditary disorders^48–51^. However, due to dependency of the detection on gene expression and relatively high noise levels, RNA-seq has not been the standard for somatic mutation detection. We demonstrated that the performance expectancy can be set by estimating the fraction of targets that are covered sufficiently for reliable detection of variants at a given allele frequency. We demonstrated that targets covered by at least 200 molecules display good performance down to an allele fraction of 5%, as most of the noise is found at the level of 3% and below^18^. Targets with lower coverage provide acceptable means of detection at higher VAF values. Target coverage depends on the tested tissue and the molecular diversity, and we demonstrated that it is closely correlated with the control gene coverage. Finally, variants of alleles that are not expressed are not subject to detection by RNA-seq.

RNA-based methods are constantly gaining space in cancer diagnostics. Parallel bulk and single-cell RNA sequencing^3^ is bringing great potential. Further efforts are required to increase awareness about factors affecting assay performance, set expectations, and allow for reliable interpretation of the results. Our work makes a significant step in this direction by providing guidance on evaluating data obtained with capture-based targeted RNA sequencing.

## ACKNOWLEDGEMENTS

We thank prof. Roger Lacave for his contribution to this work. We thank Pascal-Antoine Christin for providing critical feedback on the manuscript and Jean-Baptiste Mignardot for his contribution to visual representation of the results.

## REFERENCES

1. Oesper L, Mahmoody A, Raphael BJ. THetA: inferring intra-tumor heterogeneity from high-throughput DNA sequencing data. Genome Biol. 2013;14(7):R80. doi:10.1186/gb-2013-14-7-r80

2. Qian Y, Daza J, Itzel T, et al. Prognostic Cancer Gene Expression Signatures: Current Status and Challenges. Cells. 2021;10(3):648. doi:10.3390/cells10030648

3. Chen J, Wang X, Ma A, et al. Deep transfer learning of cancer drug responses by integrating bulk and single-cell RNA-seq data. Nat Commun. 2022;13(1):6494. doi:10.1038/s41467-022-34277-7

4. Eales JM, Jiang X, Xu X, et al. Uncovering genetic mechanisms of hypertension through multi-omic analysis of the kidney. Nat Genet. 2021;53(5):630–637. doi:10.1038/s41588-021-00835-w

5. Dorney R, Dhungel BP, Rasko JEJ, Hebbard L, Schmitz U. Recent advances in cancer fusion transcript detection. Brief Bioinform. 2023;24(1):bbac519. doi:10.1093/bib/bbac519

6. Kong-Beltran M, Seshagiri S, Zha J, et al. Somatic Mutations Lead to an Oncogenic Deletion of Met in Lung Cancer. Cancer Res. 2006;66(1):283–289. doi:10.1158/0008-5472.CAN-05-2749

7. Mertens F, Johansson B, Fioretos T, Mitelman F. The emerging complexity of gene fusions in cancer. Nat Rev Cancer. 2015;15(6):371–381. doi:10.1038/nrc3947

8. Schram AM, Chang MT, Jonsson P, Drilon A. Fusions in solid tumours: diagnostic strategies, targeted therapy, and acquired resistance. Nat Rev Clin Oncol. 2017;14(12):735–748. doi:10.1038/nrclinonc.2017.127

9. Dong Y, Xu J, Sun B, Wang J, Wang Z. MET-Targeted Therapies and Clinical Outcomes: A Systematic Literature Review. Mol Diagn Ther. 2022;26(2):203–227. doi:10.1007/s40291-021-00568-w

10. Chen J, Xu C, Lv J, et al. Clinical characteristics and targeted therapy of different gene fusions in non-small cell lung cancer: a narrative review. Transl Lung Cancer Res. 2023;12(4):895–908. doi:10.21037/tlcr-22-566

11. Wang T, Wei L, Lu Q, et al. Landscape of potentially targetable receptor tyrosine kinase fusions in diverse cancers by DNA-based profiling. Npj Precis Oncol. 2022;6(1):84. doi:10.1038/s41698-022-00325-0

12. Piskol R, Ramaswami G, Li JB. Reliable Identification of Genomic Variants from RNA-Seq Data. Am J Hum Genet. 2013;93(4):641–651. doi:10.1016/j.ajhg.2013.08.008

13. The Cancer Genome Atlas Network. Comprehensive molecular portraits of human breast tumours. Nature. 2012;490(7418):61–70. doi:10.1038/nature11412

14. Castel SE, Levy-Moonshine A, Mohammadi P, Banks E, Lappalainen T. Tools and best practices for data processing in allelic expression analysis. Genome Biol. 2015;16(1):195. doi:10.1186/s13059-015-0762-6

15. Von Ahlfen S, Missel A, Bendrat K, Schlumpberger M. Determinants of RNA Quality from FFPE Samples. Fraser P, ed. PLoS ONE. 2007;2(12):e1261. doi:10.1371/journal.pone.0001261

16. Evers DL, He J, Kim YH, Mason JT, O’Leary TJ. Paraffin Embedding Contributes to RNA Aggregation, Reduced RNA Yield, and Low RNA Quality. J Mol Diagn. 2011;13(6):687–694. doi:10.1016/j.jmoldx.2011.06.007

17. Hedegaard J, Thorsen K, Lund MK, et al. Next-Generation Sequencing of RNA and DNA Isolated from Paired Fresh-Frozen and Formalin-Fixed Paraffin-Embedded Samples of Human Cancer and Normal Tissue. Zuo Z, ed. PLoS ONE. 2014;9(5):e98187. doi:10.1371/journal.pone.0098187

18. Graw S, Meier R, Minn K, et al. Robust gene expression and mutation analyses of RNA-sequencing of formalin-fixed diagnostic tumor samples. Sci Rep. 2015;5(1):12335. doi:10.1038/srep12335

19. Wehmas LC, Wood CE, Chorley BN, Yauk CL, Nelson GM, Hester SD. Enhanced Quality Metrics for Assessing RNA Derived From Archival Formalin-Fixed Paraffin-Embedded Tissue Samples. Toxicol Sci. 2019;170(2):357–373. doi:10.1093/toxsci/kfz113

20. Verwilt J, Mestdagh P, Vandesompele J. Artifacts and biases of the reverse transcription reaction in RNA sequencing. RNA. 2023;29(7):889–897. doi:10.1261/rna.079623.123

21. Minshall N, Git A. Enzyme- and gene-specific biases in reverse transcription of RNA raise concerns for evaluating gene expression. Sci Rep. 2020;10(1):8151. doi:10.1038/s41598-020-65005-0

22. Guo Y, Zhao S, Sheng Q, Samuels DC, Shyr Y. The discrepancy among single nucleotide variants detected by DNA and RNA high throughput sequencing data. BMC Genomics. 2017;18(S6):690. doi:10.1186/s12864-017-4022-x

23. Zhang P, Samuels DC, Lehmann B, et al. Mitochondria sequence mapping strategies and practicability of mitochondria variant detection from exome and RNA sequencing data. Brief Bioinform. 2016;17(2):224–232. doi:10.1093/bib/bbv057

24. Buckley J, Schmidt R, Ostrow D, et al. An Exome Capture-Based RNAseq Assay for Genome-Wide Identification and Prioritization of Clinically Important Fusions in Pediatric Tumors. J Mol Diagn. Published online November 2023:S152515782300274X. doi:10.1016/j.jmoldx.2023.11.003

25. Babiceanu M, Qin F, Xie Z, et al. Recurrent chimeric fusion RNAs in non-cancer tissues and cells. Nucleic Acids Res. 2016;44(6):2859–2872. doi:10.1093/nar/gkw032

26. Li X, Zhang P, Wang H, Yu Y. Genes expressed at low levels raise false discovery rates in RNA samples contaminated with genomic DNA. BMC Genomics. 2022;23(1):554. doi:10.1186/s12864-022-08785-1

27. Dobin A, Davis CA, Schlesinger F, et al. STAR: ultrafast universal RNA-seq aligner. Bioinformatics. 2013;29(1):15–21. doi:10.1093/bioinformatics/bts635

28. Li H, Durbin R. Fast and accurate short read alignment with Burrows–Wheeler transform. Bioinformatics. 2009;25(14):1754–1760. doi:10.1093/bioinformatics/btp324

29. Sheng Q, Vickers K, Zhao S, et al. Multi-perspective quality control of Illumina RNA sequencing data analysis. Brief Funct Genomics. Published online September 29, 2016:elw035. doi:10.1093/bfgp/elw035

30. Kumar G, Ertel A, Feldman G, Kupper J, Fortina P. iSeqQC: a tool for expression-based quality control in RNA sequencing. BMC Bioinformatics. 2020;21(1):56. doi:10.1186/s12859-020-3399-8

31. Williams AG, Thomas S, Wyman SK, Holloway AK. RNA-seq Data: Challenges in and Recommendations for Experimental Design and Analysis. Curr Protoc Hum Genet. 2014;83(1). doi:10.1002/0471142905.hg1113s83

32. Jones W, Greytak S, Odeh H, et al. Deleterious effects of formalin-fixation and delays to fixation on RNA and miRNA-Seq profiles. Sci Rep. 2019;9(1):6980. doi:10.1038/s41598-019-43282-8

33. Adiconis X, Borges-Rivera D, Satija R, et al. Comparative analysis of RNA sequencing methods for degraded or low-input samples. Nat Methods. 2013;10(7):623–629. doi:10.1038/nmeth.2483

34. Barua S, Wang G, Mansukhani M, Hsiao S, Fernandes H. Key considerations for comprehensive validation of an RNA fusion NGS panel. Pract Lab Med. 2020;21:e00173. doi:10.1016/j.plabm.2020.e00173

35. Reeser JW, Martin D, Miya J, et al. Validation of a Targeted RNA Sequencing Assay for Kinase Fusion Detection in Solid Tumors. J Mol Diagn. 2017;19(5):682–696. doi:10.1016/j.jmoldx.2017.05.006

36. Desmeules P, Boudreau DK, Bastien N, et al. Performance of an RNA-Based Next-Generation Sequencing Assay for Combined Detection of Clinically Actionable Fusions and Hotspot Mutations in NSCLC. JTO Clin Res Rep. 2022;3(2):100276. doi:10.1016/j.jtocrr.2022.100276

37. Kaya C, Dorsaint P, Mercurio S, et al. Limitations of Detecting Genetic Variants from the RNA Sequencing Data in Tissue and Fine-Needle Aspiration Samples. Thyroid. 2021;31(4):589–595. doi:10.1089/thy.2020.0307

38. Fu GK, Xu W, Wilhelmy J, et al. Molecular indexing enables quantitative targeted RNA sequencing and reveals poor efficiencies in standard library preparations. Proc Natl Acad Sci. 2014;111(5):1891–1896. doi:10.1073/pnas.1323732111

39. Matsubara T, Soh J, Morita M, et al. DV200 Index for Assessing RNA Integrity in Next-Generation Sequencing. BioMed Res Int. 2020;2020:1–6. doi:10.1155/2020/9349132

40. Norton N, Sun Z, Asmann YW, et al. Gene Expression, Single Nucleotide Variant and Fusion Transcript Discovery in Archival Material from Breast Tumors. Dadras SS, ed. PLoS ONE. 2013;8(11):e81925. doi:10.1371/journal.pone.0081925

41. Jennings LJ, Arcila ME, Corless C, et al. Guidelines for Validation of Next-Generation Sequencing–Based Oncology Panels. J Mol Diagn. 2017;19(3):341–365. doi:10.1016/j.jmoldx.2017.01.011

42. Heydt C, Wölwer CB, Velazquez Camacho O, et al. Detection of gene fusions using targeted next-generation sequencing: a comparative evaluation. BMC Med Genomics. 2021;14(1):62. doi:10.1186/s12920-021-00909-y

43. Zhao Y, Mehta M, Walton A, et al. Robustness of RNA sequencing on older formalin-fixed paraffin-embedded tissue from high-grade ovarian serous adenocarcinomas. Real FX, ed. PLOS ONE. 2019;14(5):e0216050. doi:10.1371/journal.pone.0216050

44. Bieler J, Kubik S, Macheret M, Pozzorini C, Willig A, Xu Z. Benefits of applying molecular barcoding systems are not uniform across different genomic applications. J Transl Med. 2023;21(1):305. doi:10.1186/s12967-023-04160-0

45. Fu C, Marczyk M, Samuels M, et al. Targeted RNAseq assay incorporating unique molecular identifiers for improved quantification of gene expression signatures and transcribed mutation fraction in fixed tumor samples. BMC Cancer. 2021;21(1):114. doi:10.1186/s12885-021-07814-8

46. Peng H, Huang R, Wang K, et al. Development and Validation of an RNA Sequencing Assay for Gene Fusion Detection in Formalin-Fixed, Paraffin-Embedded Tumors. J Mol Diagn. 2021;23(2):223–233. doi:10.1016/j.jmoldx.2020.11.005

47. Park HJ, Baek I, Cheang G, Solomon JP, Song W. Comparison of RNA-Based Next-Generation Sequencing Assays for the Detection of NTRK Gene Fusions. J Mol Diagn. 2021;23(11):1443–1451. doi:10.1016/j.jmoldx.2021.07.027

48. Shah SP, Köbel M, Senz J, et al. Mutation of FOXL2 in Granulosa-Cell Tumors of the Ovary. N Engl J Med. 2009;360(26):2719–2729. doi:10.1056/NEJMoa0902542

49. Kridel R, Meissner B, Rogic S, et al. Whole transcriptome sequencing reveals recurrent NOTCH1 mutations in mantle cell lymphoma. Blood. 2012;119(9):1963–1971. doi:10.1182/blood-2011-11-391474

50. Chandrasekharappa SC, Lach FP, Kimble DC, et al. Massively parallel sequencing, aCGH, and RNA-Seq technologies provide a comprehensive molecular diagnosis of Fanconi anemia. Blood. 2013;121(22):e138–e148. doi:10.1182/blood-2012-12-474585

51. Martínez-Ruiz C, Black JRM, Puttick C, et al. Genomic–transcriptomic evolution in lung cancer and metastasis. Nature. 2023;616(7957):543–552. doi:10.1038/s41586-023-05706-4

