## Supplemental Figures for "Critical assay parameters facilitating confident detection of expression changes, fusions and short variants in RNA isolated from tissue"

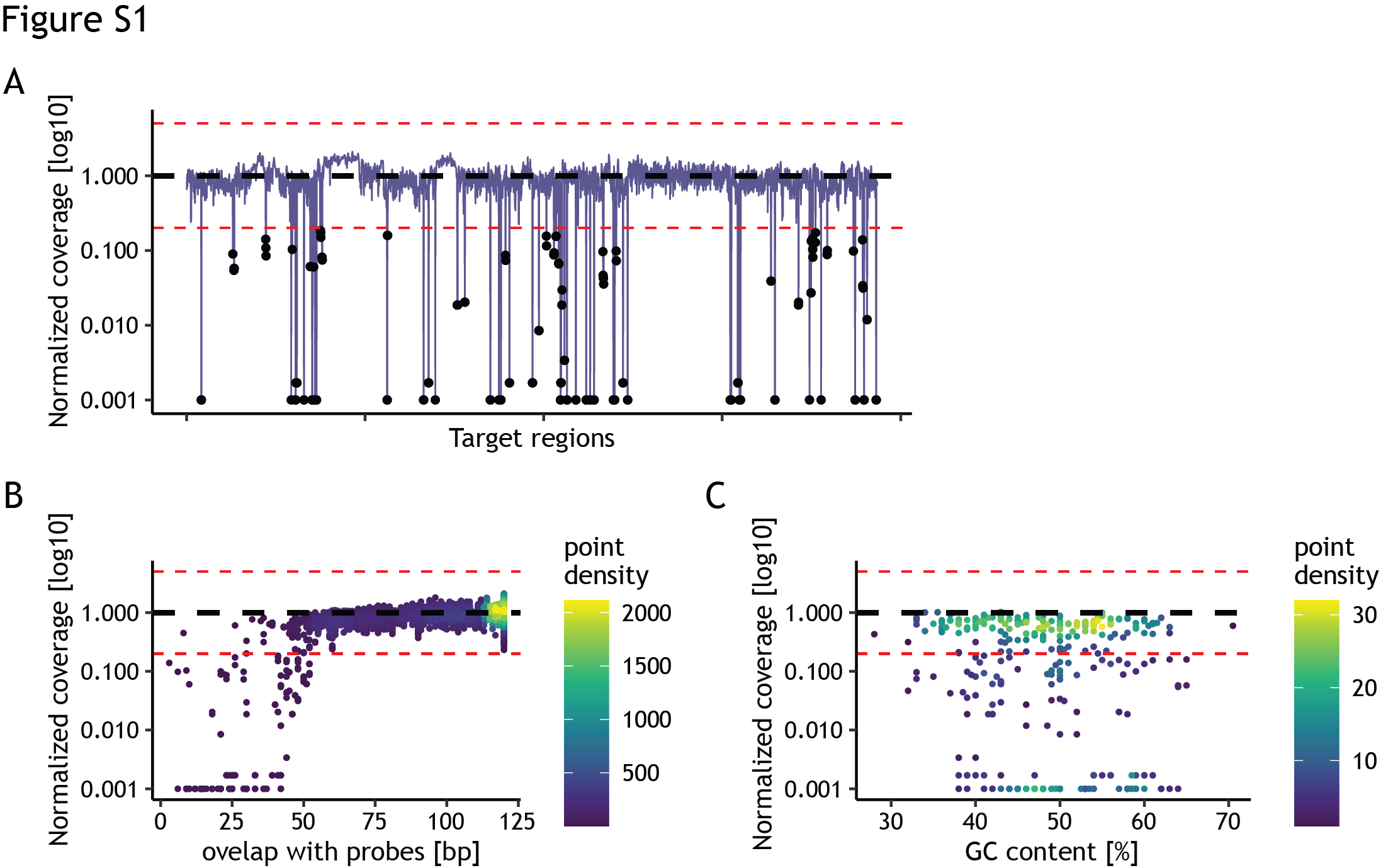

**Figure S1. (A)** Median-normalized coverage for all panel regions captured using DNA sample as input; red lines delineate 1/5 and 5x of the median. **(B)** Median-normalized coverage for all panel regions, using DNA sample as input, as a function of length of the total probe overlap of that region in DNA. **(C)** Median-normalized coverage for all panel regions, using DNA sample as input, as a function of GC content of the target region.

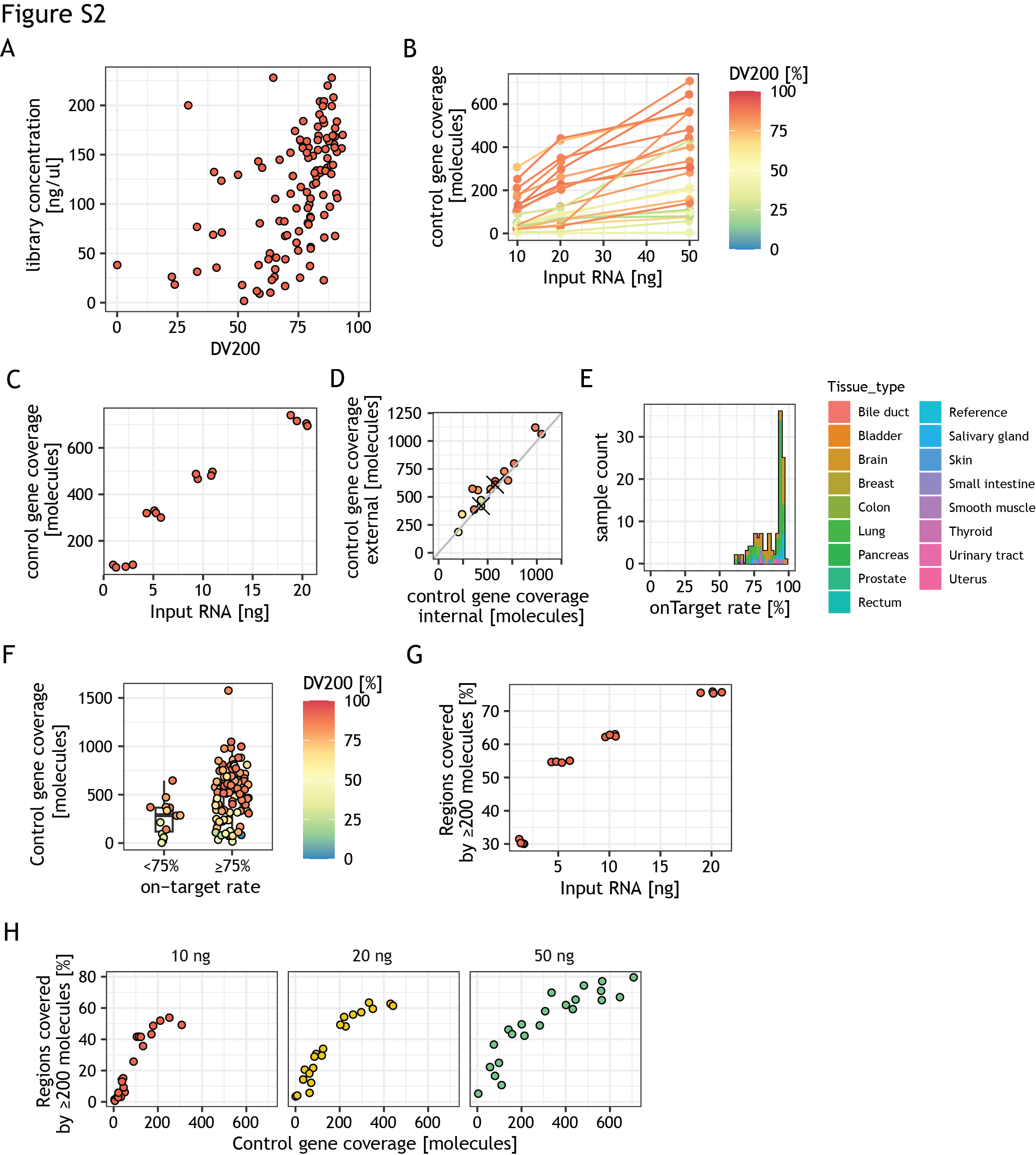

**Figure S2.** **(A)** Relationship between DV200 value (x axis) and obtained library concentration (Y axis) for all samples for which 50 ng RNA was processed. **(B)** Control gene coverage (y axis) measured for samples for which 3 different input quantities (x axis) were processed. **(C)** Control gene coverage (y axis) measured for replicates of a single sample for which different input quantities (x axis) were processed. **(D)** Control gene coverage measured for 13 clinical samples in the internal laboratory (x axis) and external laboratory (y axis); two samples processed in duplicate are marked with a cross. **(E)** Distribution of samples depending on their fraction of reads falling within the panel targets (x axis); color indicates sample source tissue. **(F)** Control gene coverage depending on sample on-target rate being at elast 75% or below. **(G)** Fraction of target regions covered by at least 200 molecules (y axis) in replicates of a sample for which different input quantities (x axis) were processed. **(H)** As for G but for all clinical samples for which 3 different input quantities (10, 20 and 50 ng, left to right, respectively) were processed.

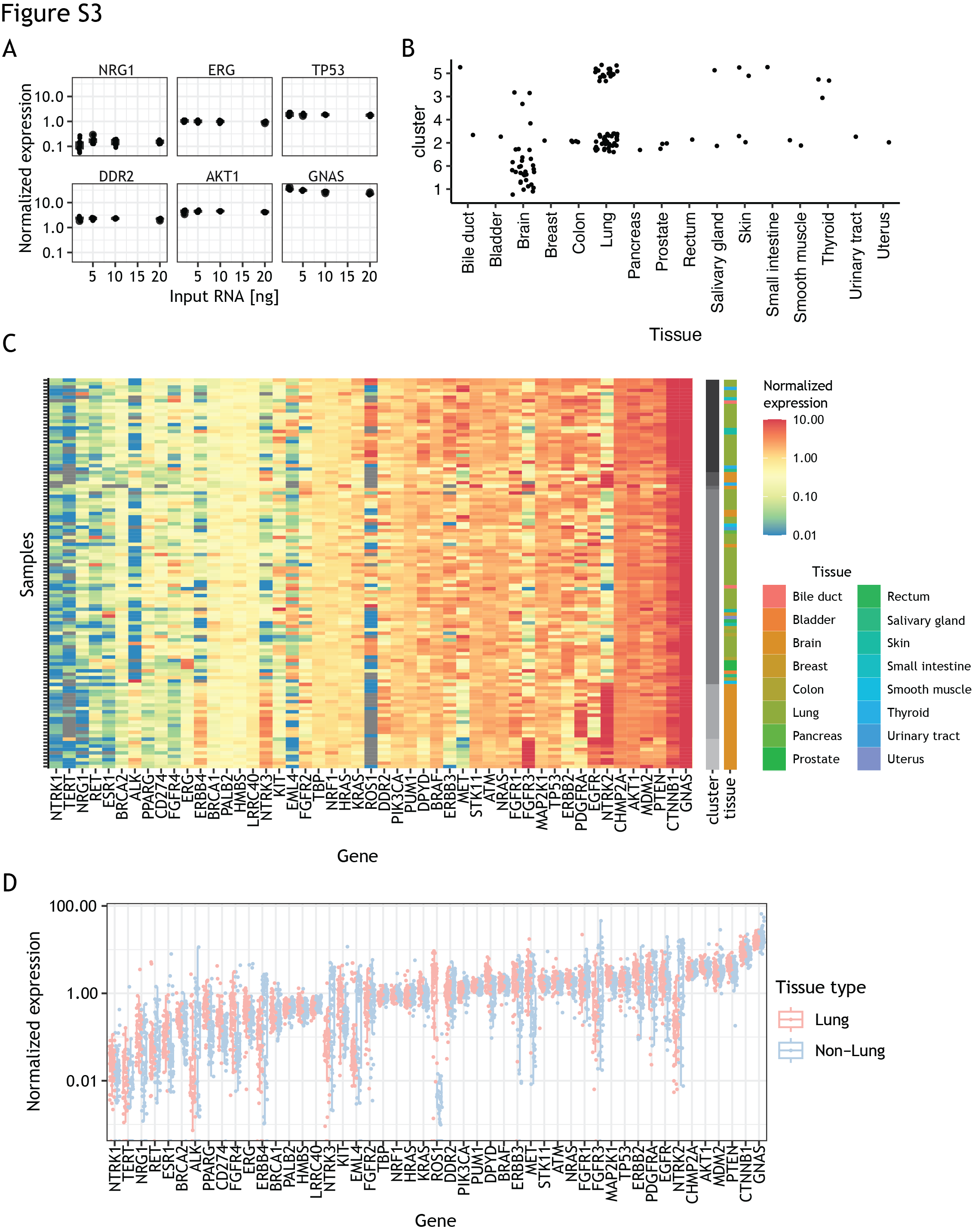

**Figure S3. (A)** Normalized expression values for 6 representative genes measured in replicates of sample for which different input quantities (x axis) were processed. **(B)** Sample clusters obtained by k-means clustering (k=6) using measured expression values. **(C)** Heatmap of expression values for all samples arranged according to the k-means clustering; sample tissue displayed as color on the right. **(D)** Normalized expression values (y axis) measured for all genes included in the panel and colored depending on the source tissue for lung (red) and non-lung (red).

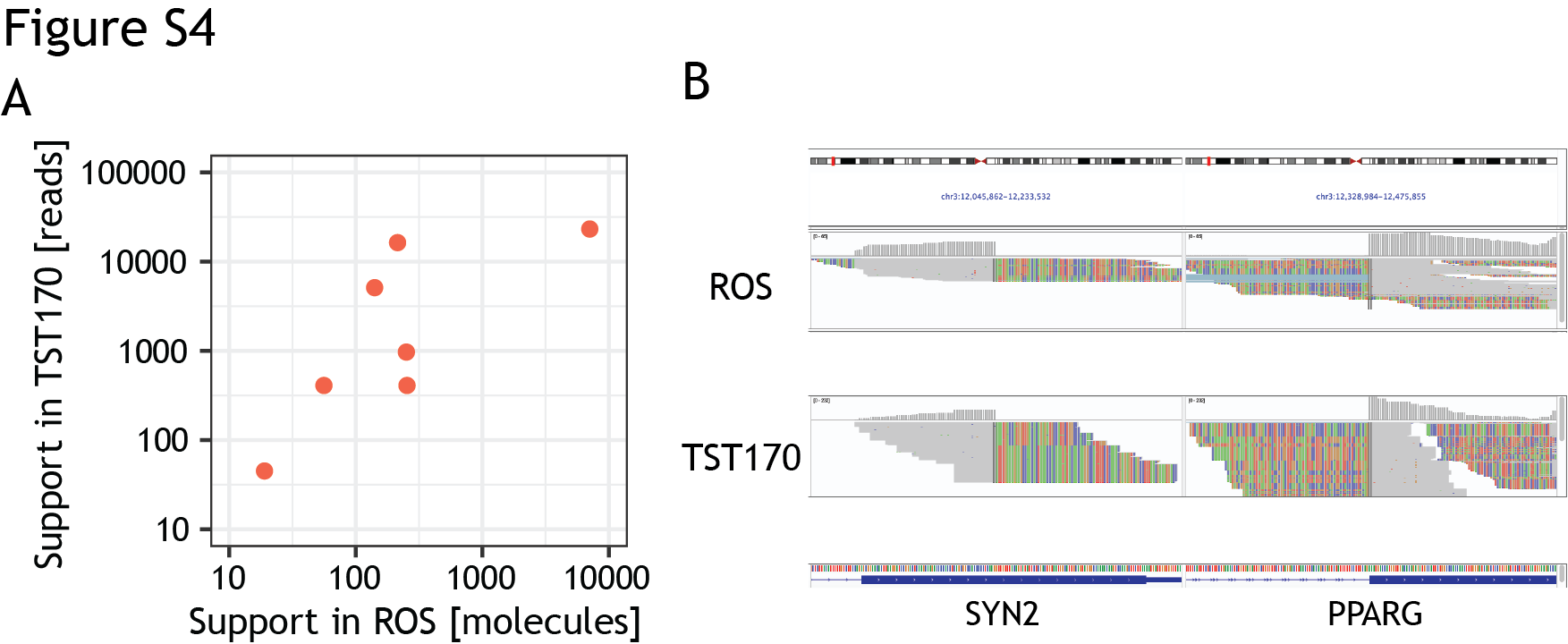

**Figure S4. (A)** Fusion-supporting molecule counts determined with ROS (x axis) and read count for these events measured with TST170 (y axis) for the same subset of samples processed with both methods. **(B)** Snapshots from the genome browser of reads supporting a SYN2-PPARG fusion, bam files derived from analyses with ROS (top) and TST170 (bottom); the fusion was not reported by TST170.

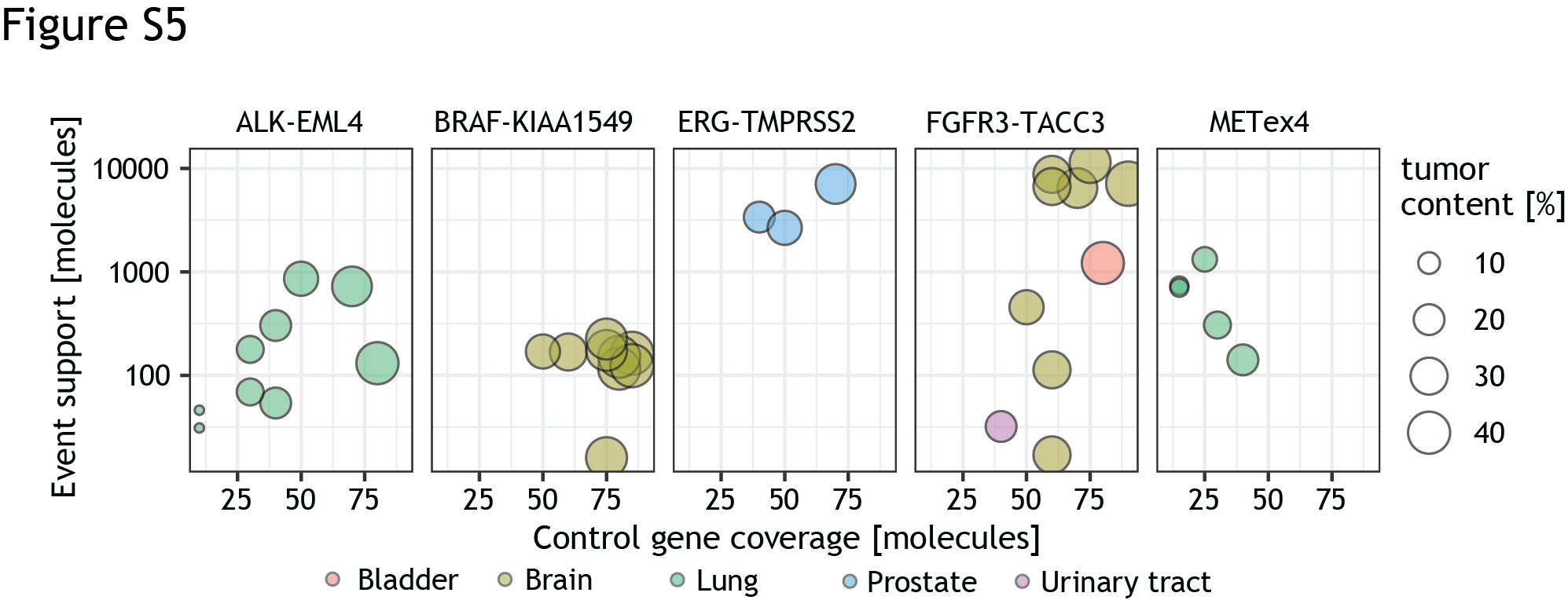

**Figure S5.** Supporting molecule count (y axis) as a function of tumor content (x axis) for events present in at least three clinical samples; points are colored according to source tissue.

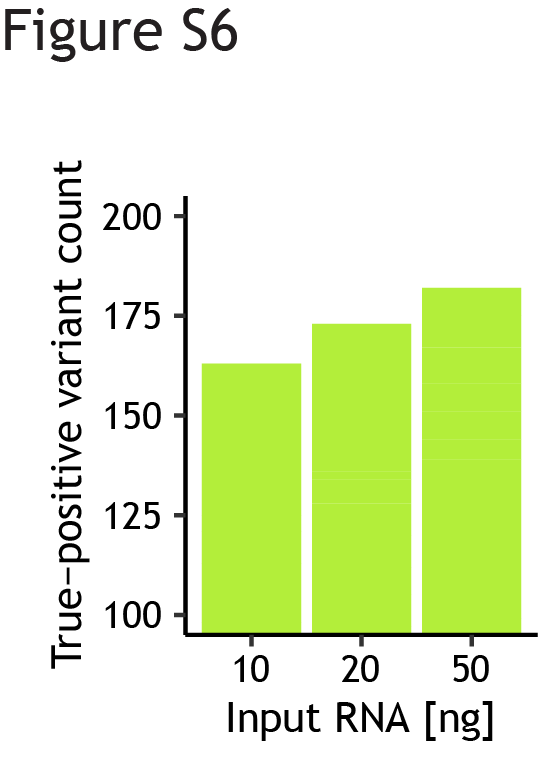

**Figure S6.** Total count of true-positive variants depending on the input quantity for samples for which 3 different input quantities were processed.

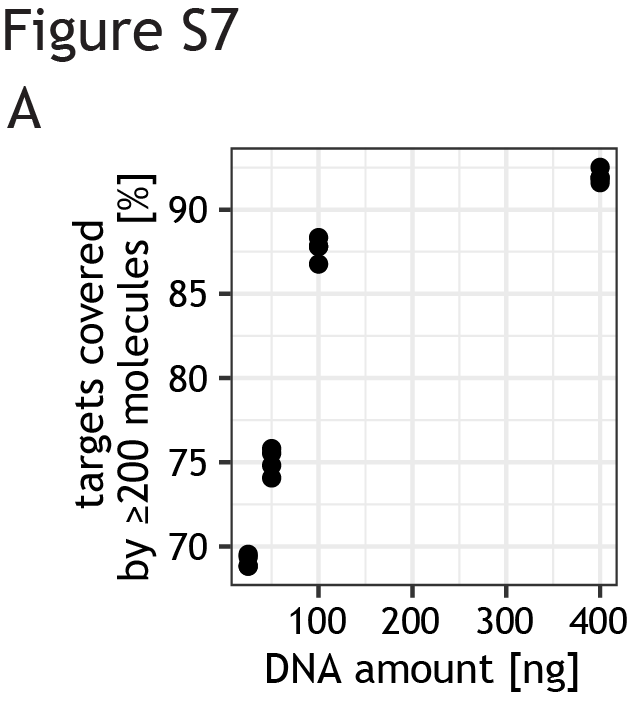

**Figure S7.** Fraction of target regions covered by at least 200 molecules (y axis) in replicates of a sample for which 50 ng of RNA was mixed with different DNA quantities (x axis).
