## Supplemental Methods for "Critical assay parameters facilitating confident detection of expression changes, fusions and short variants in RNA isolated from tissue"

**Concordance of fusion detection**

To evaluate the performance of detection of RNA-level rearrangements, information about fusions and exon skipping events detected in the samples processed with ROS were compared to prior results obtained using a widely used nested amplicon-based approach (Archer FUSIONPlex). Additionally, as a control, a reference sample bearing 18 events targeted by ROS was used in duplicate.

After initial concordance analysis, for samples with discordant results, an additional test was performed using another hybrid capture-based method, TST170, previously reported to achieve the best sensitivity among several assays^27^. For one sample, the remaining material was insufficient to repeat the test. The results of the TST170 assay were considered as the ground truth for discordant samples (**Supplemental Table 2**).

One of the two false-positive events, ROS1-PCMDT1, was supported by only 10 molecules, the applied threshold value for detection. Moreover, the sample had tumor content of only 10% and relatively poor CGC (75 molecules). On the contrary, the SYN2-PPARG fusion was detected in good quality sample (CGC of 1048 molecules), was supported by 24 molecules in ROS data and fusion-supporting reads were also observed in TST170 results despite the event not being reported by the assay (**Figure S4B**). Curiously, the breakpoint on SYN2 was found in the exon. These results may indicate the presence of low levels of a chimeric transcript.
